# RNA Polymerase II, the BAF remodeler and transcription factors synergize to evict nucleosomes

**DOI:** 10.1101/2023.01.22.525083

**Authors:** Sandipan Brahma, Steven Henikoff

## Abstract

Chromatin accessibility is a hallmark of active transcription and requires ATP-dependent nucleosome remodeling by Brahma-Associated Factor (BAF). However, the mechanistic link between transcription, nucleosome remodeling, and chromatin accessibility is unclear. Here, we used a chemical-genetic approach to dissect the interplay between RNA Polymerase II (RNAPII), BAF, and DNA-sequence-specific transcription factors (TFs) in mouse embryonic stem cells. By time-resolved chromatin profiling with acute transcription block at distinct stages, we show that RNAPII promoter-proximal pausing stabilizes BAF chromatin occupancy and enhances nucleosome eviction by BAF. We find that RNAPII and BAF probe both transcriptionally active and Polycomb-repressed genomic regions and provide evidence that TFs capture transient site exposure due to nucleosome unwrapping by BAF to confer locus specificity for persistent chromatin remodeling. Our study reveals the mechanistic basis of cell-type-specific chromatin accessibility. We propose a new paradigm for how functional synergy between dynamically acting chromatin factors regulates nucleosome organization.

## Introduction

The positions of nucleosomes with respect to gene-regulatory elements such as promoters and enhancers is fundamental to transcription regulation. Nucleosomes occlude DNA sequences from transcription factors (TF) and prevent loading of the basal transcription machinery including RNA Polymerase II (RNAPII)^1,2^. Regulatory elements of transcriptionally active genes are generally considered to lack positioned nucleosomes and are therefore accessible to protein factors^3^. Identification of such nucleosome depleted regions (NDRs) is a standard practice for inferring transcription activity^4^. However, several recent studies have shown that even at the steady state, promoters and enhancers of transcriptionally active genes are not stably depleted of nucleosomes^5–8^. Rather, a dynamic cycle of nucleosome loading and eviction characterizes transcriptionally active chromatin such that genomic regions are never completely accessible or occluded in a population of cells^5,6,9^.

In general, the causal relationship between chromatin accessibility and transcription remains unclear^6,10,11^. The most widely accepted model posits that a special class of “pioneer” TFs that can bind to partial DNA sequence motifs wrapped around nucleosomal histones recruit transcriptional activators such as ATP-dependent nucleosome remodelers. These factors then evict nucleosomes to facilitate the binding of secondary TFs and RNAPII^12,13^. Although parts of this hierarchical model have been tested by *in vitro* experiments, how factors functionally cooperate *in vivo* remains unclear^8,14–16^. Genome-wide analysis of RNAPII and nucleosome occupancy in *Drosophila* have shown that promoter-proximal pausing of RNAPII counteracts promoter nucleosome occupancy, and therefore, may facilitate NDR formation^17,18^. In mammalian cells, accessibility of promoters and enhancers is consistently associated with paused RNAPII downstream of the NDR^19,20^. These studies suggest a role of RNAPII pausing in promoter and enhancer nucleosome eviction. RNAPII pausing was first described at the *Drosophila* heat shock genes, where RNAPII synthesizes the first 25-50 nucleotides of RNA and then pauses, waiting for an activating signal^21^. Heat shock rapidly triggered the release of paused RNAPII into the gene body for a robust transcriptional response^22^. Although decades of work have established RNAPII pausing as a common regulatory strategy in higher eukaryotes^23^, its functional role still remains unclear.

We hypothesized that RNAPII pausing may facilitate ATP-dependent chromatin remodeling to form NDRs. ATP-dependent chromatin remodelers are multi-subunit protein complexes belonging to one of four main families (SWI/ SNF, ISWI, INO80, and CHD), which dynamically regulate nucleosome positions^24^. In animals, the main remodelers evicting nucleosomes from gene promoters and enhancers are the SWI/SNF (SWItch independent/ Sucrose Non-Fermenting) family remodelers, for example, the mammalian BAF (Brahma-associated Factor) complex^9^. BAF complexes consist of the catalytic subunit BRG1 (Brahma-related gene 1) and 15-20 additional subunits, most of which are evolutionarily conserved. Consistent with its fundamental role in regulating nucleosome organization, BAF is essential for almost all developmental gene regulation, and BAF subunits are recurrently mutated in more than 20% of all human cancers^25^. We have previously found that the homologous complex in *Saccharomyces cerevisiae*, RSC (Remodeling the Structure of Chromatin), is associated with partially unwrapped nucleosomal intermediates at transcriptionally active gene promoters^5^. We further showed that the general-regulatory TFs (GRFs) Abf1 and Reb1 bind their cognate sequence motifs within these partially unwrapped nucleosomes, suggesting that a dynamic cycle of nucleosome formation and depletion characterizes transcriptionally active promoters^5,26^. Consistent with our cyclical model, it was recently shown in mouse embryonic stem cells (mESCs) and human cancer cell lines that ATP-dependent nucleosome remodeling by BAF is continuously required throughout the cell cycle to maintain nucleosome depletion and DNA accessibility at promoters and enhancers^9,27^. However, the possible role of RNAPII in these dynamic processes has not been explored.

In this study, we used highly specific small molecule inhibitors to block RNAPII at either transcription initiation or release from promoter-proximal pausing to determine drug-induced kinetic changes in RNAPII, BAF and nucleosome occupancy in mESCs. We show that RNAPII promoter-proximal pausing promotes BAF chromatin binding and remodeling, leading to enhanced nucleosome eviction and DNA accessibility at gene promoters and distal regulatory regions. Although RNAPII and BAF engage chromatin genome wide, effective chromatin remodeling occurs only at active regulatory elements and requires a tripartite synergy between RNAPII, BAF, and DNA-sequence-specific TFs for nucleosome eviction. Our study broadly explains how modulating the dynamics of chromatin factors can drive altered chromatin structure and gene expression, such as in development and cancer.

## Results

### BAF chromatin occupancy is strongly dependent on RNAPII

We used the Cleavage Under Targets and Tagmentation (CUT&Tag) method to determine the occupancy of RNAPII and BAF in chromatin from mESCs. In CUT&Tag, antibodies targeting chromatin-associated proteins are added to permeabilized cells, followed by a proteinA-Tn5 transposasome fusion, which binds to the antibody-targeted chromatin regions. Upon activation, the tethered Tn5 inserts DNA-sequencing adapters into targeted genomic regions, allowing us to map protein binding genome-wide with high specificity and resolution^28^. For RNAPII, we used antibodies targeting the heptapeptide repeat within the C-terminal domain ofthe largest subunit, RPB1, phosphorylated at serine 5 (RNAPII-S5P) or serine 2 (RNAPII-S2P), and another core RNAPII subunit, RPB3. CUT&Tag showed strong occupancy of RNAPII near the transcription start sites (TSSs) of genes within promoters, as well as at promoter-distal regions corresponding to annotated transcriptional enhancers (Extended Data Fig. 1a-d). RNAPII-occupied sites showed strong enrichment for histones containing posttranslational modifications characteristic of active transcription^11^, such as histone H3 with mono- or tri-methylation of lysine 4 (H3K4me1 and H3K4me3), which mark enhancers and promoters respectively. In contrast, promoters enriched for histone H3 with the repressive trimethylation of lysine 27 (H3K27me3) showed little RNAPII occupancy (Extended Data Fig. 1a-c). To determine the occupancy of the BAF complex, we applied CUT&Tag by targeting the BRG1 subunit. BRG1 includes the catalytic domains responsible for ATP hydrolysis and DNA translocation, essential for nucleosome remodeling^29^. Although BAF complexes may contain the alternative BRM ATPase subunits, BRM is not expressed in mESCs, therefore BRG1 represents all functional BAF complexes in these cells. CUT&Tag showed that BAF occupies the same genomic regions as RNAPII and showed strong association with promoter-proximal pausing of RNAPII, characterized by serine 5 phosphorylation of RPB1 (RNAPII-S5P) (Fig. 1a, Extended Data Fig. 1a-e). We have recently demonstrated that low-salt tagmentation conditions for RNAPII-S5P CUT&Tag produce high-resolution maps of transcription-coupled accessible regulatory sites including active gene promoters and enhancers^19,20^. RNAPII-S5P-associated accessible chromatin regions in mESCs, hereby referred to as S5P CUTAC (Cleavage Under Targeted Accessible Chromatin) peaks mapped within NDRs determined by micrococcal nuclease (MNase) digestion of chromatin, and immediately upstream of nascent RNA transcription start sites determined by START-seq^30,31^ (Extended Data Fig. 1a,f). Therefore, S5P CUTAC peaks represent NDRs upstream of genic promoter TSSs and start sites of enhancer RNA transcription, selectively in cells where these loci are occupied by RNAPII. When aligned over S5P CUTAC peaks, BRG1 CUT&Tag showed strong BAF occupancy, consistent with the function of BAF in generating and/or maintaining the NDRs (Fig. 1a, Extended Data Fig. 1a,b).

**Fig. 1.**
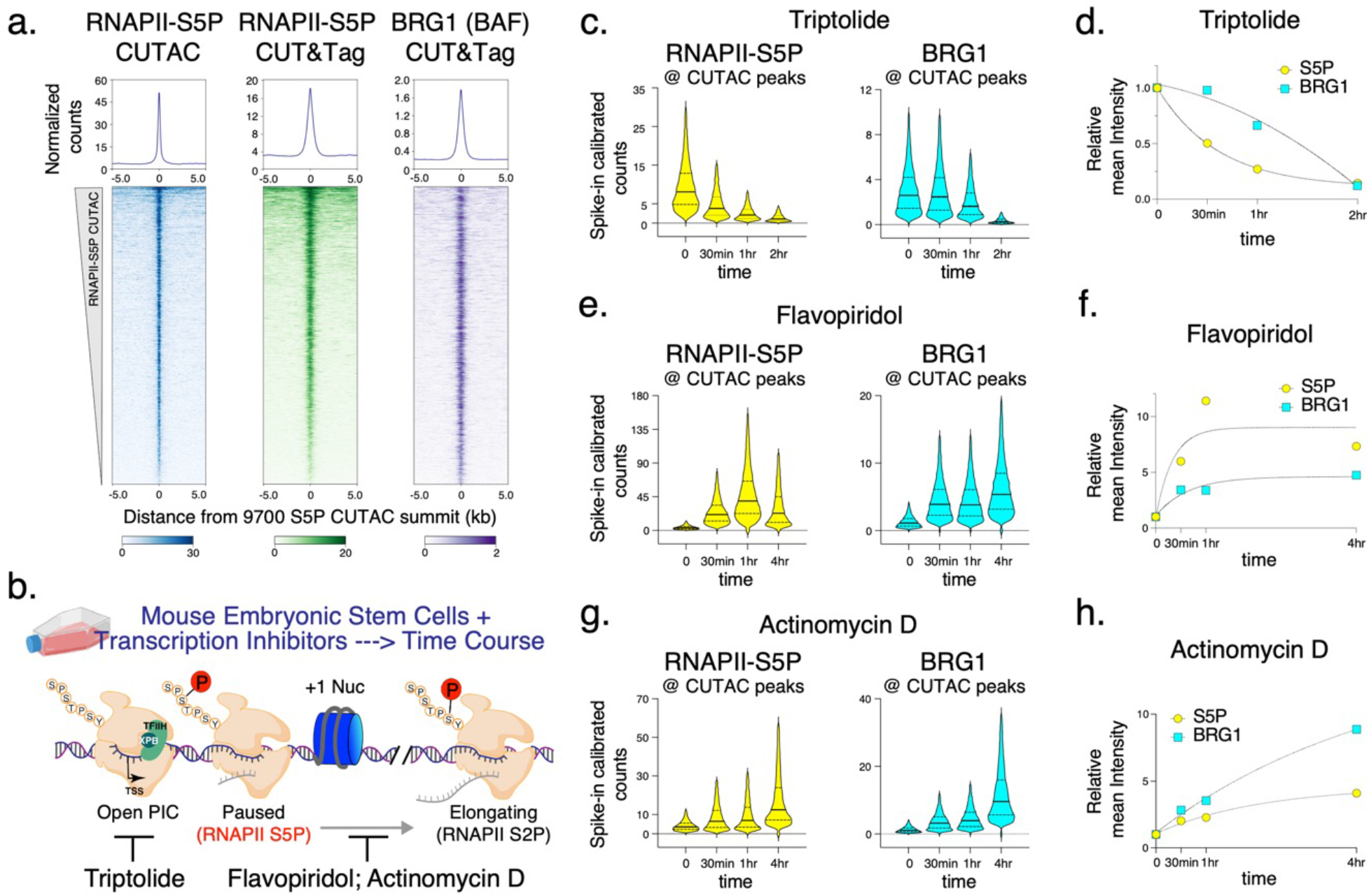
RNAPII promoter proximal pausing facilitates BAF chromatin binding. **a,** Heatmaps (bottom) and average plots (top) comparing chromatin accessibility assayed by S5P CUTAC, with RNAPII-S5P and BRG1 occupancy (CUT&Tag), relative to the primary peaks (summits) of S5P CUTAC and sorted by decreasing accessibility (CUTAC signal). **b,** Schematic showing distinct stages in RNAPII transcription that are inhibited by drugs used in this study. **c, e, g,** Violin plots of spike-in calibrated CUT&Tag signal distribution comparing RNAPII-S5P and BRG1 occupancy over S5P CUTAC peaks at time points after drug treatments for transcription inhibition. Median value (solid line), upper- and lower quartiles (broken lines) and outliers were calculated using the Tukey method. **d, f, h,** Fold changes in mean RNAPII-S5P and BRG1 occupancy (spike-in calibrated CUT&Tag) over S5P CUTAC peaks at time points after drug treatments. Datasets are representative of at least two biological replicates.

To determine whether and how RNAPII regulates BAF, we applied a chemical-genetic approach using fast-acting cell-permeable small molecule inhibitors to acutely inhibit transcription at distinct stages and modulate RNAPII dynamics (Fig. 1b). We selected three inhibitors, Triptolide, Flavopiridol and Actinomycin D, which affect the RNAPII transcription cycle in specific and distinct ways^32^. The natural diterpene triepoxide Triptolide inhibits ATPase activity of the XPB subunit of TFIIH, which is a part of the transcription pre-initiation complex (PIC) consisting of RNAPII and several additional cofactors. Triptolide prevents transcription initiation by blocking the ATP-dependent activity of XPB to translocate DNA into the RNAPII active site and so induces a fast proteasomal degradation of the RNAPII subunit RPB1^33^. We subjected mESC cultures to a time course of treatment with 10 μM Triptolide. Although we did not observe changes in cell or colony morphology and cell viability for up to two hours, cells started to dislodge and lose viability at four hours of treatment. CUT&Tag showed a dramatic and rapid loss of RNAPII-S5P occupancy genome wide (Fig. 1c,d, Extended Data Fig. 2a). We observed a 50% reduction in occupancy within 30 min, and almost all RNAPII-S5P was lost within two hours. Here and elsewhere, we used spike-in calibration, which is vital for quantifying such genome-wide differences. CUT&Tag for BRG1 showed a similar response, that is, a rapid loss of BAF occupancy genome-wide upon Triptolide treatment (Fig. 1c,d, Extended Data Fig. 2b). This implies that RNAPII and transcription initiation (either RNAPII loading or promoter-proximal pausing) facilitates BAF chromatin binding. Interestingly, BAF was lost at a slower rate than RNAPII-S5P, suggesting that RNAPII-S5P may turn over faster than BAF (Fig. 1d).

To distinguish whether BAF binding is facilitated by RNAPII loading or pausing, we treated mESCs with inhibitors that are expected to accumulate paused RNAPII by inhibiting the transition to productive elongation. Flavopiridol is a semi-synthetic flavonoid which inhibits the kinase activity of CDK9, a component of the transcription elongation factor pTEFb, and is known to increase RNAPII pausing genome wide^32,34^. As expected, CUT&Tag mapping showed a rapid increase in RNAPII-S5P over a four-hour time course following 1 μM Flavopiridol treatment (Fig. 1e,f, Extended Data Fig. 2e). A corresponding rapid increase in BRG1 CUT&Tag established that RNAPII pausing promotes BAF chromatin occupancy. This effect is not limited to Flavopiridol, as we observed similar rapid increases in both RNAPII-S5P and BRG1 over a time course of treatment with 5 μg/mL Actinomycin D (Fig. 1g,h, Extended Data Fig. 2e,f). Actinomycin D inhibits transcription by a distinct mechanism as it intercalates within unwound DNA strands at the active site of RNAPII and acts as a physical roadblock to RNAPII elongation^32^. Live-cell single-molecule imaging has shown that Actinomycin D increases the residence time of RNAPII-S5P on chromatin, while Flavopiridol does not^35^. This implies that Actinomycin D “locks” paused RNAPII, while Flavopiridol increases the population of paused RNAPII, with rapid turnover. In the presence of Flavopiridol, CUT&Tag signal for RNAPII-S5P increased early but plateaued, while in Actinomycin D RNAPII-S5P occupancy continued to increase (Fig. 1f,h). BRG1 CUT&Tag showed the same patterns as RNAPII-S5P, which further demonstrates the strong dependence of BAF on paused RNAPII for chromatin binding (Fig. 1f-h). Although RNAPII-S5P and BRG1 showed higher occupancy at promoter-distal CUTAC sites compared to gene promoters (Extended Data Fig. 1d), treatment with the transcription inhibitors resulted in identical kinetics at both sets of regions, confirming that promotion of BAF occupancy by paused RNAPII is a general phenomenon occurring broadly across the mESC genome (Extended Data Fig. 2c,d,g-j).

### BAF unwraps and evicts nucleosomes to establish NDRs

Having found that paused RNAPII facilitates BAF chromatin binding, we next investigated its effect on nucleosome occupancy and chromatin accessibility at gene promoters and distal regulatory regions using RNAPII-S5P CUTAC. Analysis of DNA-insert sizes in CUTAC libraries can show how closely two Tn5 molecules could integrate into the same DNA, providing a protein footprint. A similar approach using untargeted Tn5 in ATAC-seq is routinely used to determine protein occupancy at regulatory regions^36,37^. RNAPII-S5P CUTAC restricts this analysis to cells where a particular genomic locus is actively transcribed or at least occupied by RNAPII. RNAPII-S5P CUTAC in mESCs produced fragments that were mostly shorter than 120 base pairs (bp) (>85% of total reads overlapping CUTAC peaks, Fig. 2a, Extended Data Fig. 3a,b; DMSO only), suggesting that transcriptionally active gene promoters and distal regulatory regions are mostly occupied by proteins with footprints smaller than nucleosomes.

**Fig. 2.**
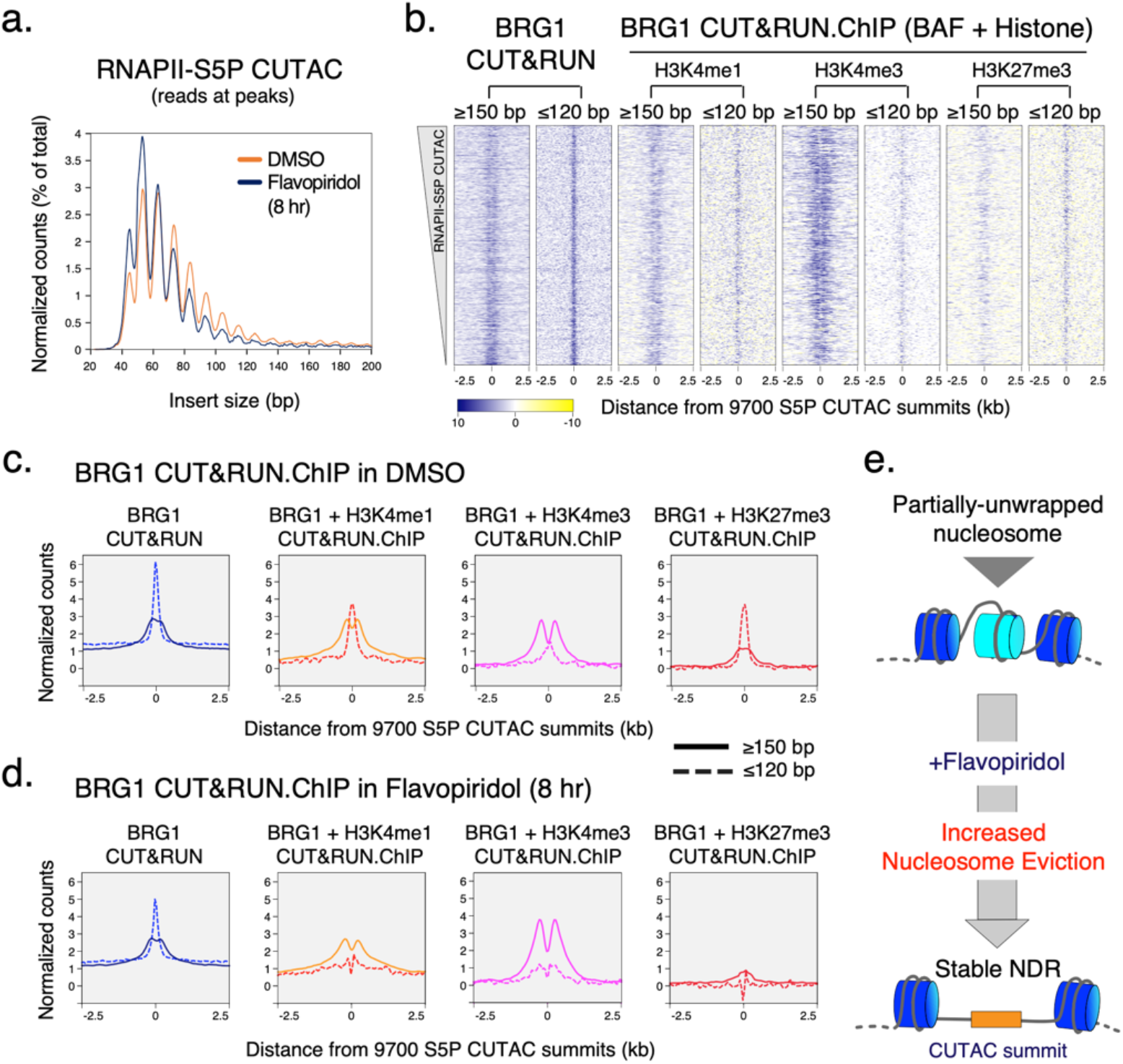
Enrichment of BAF increases nucleosome eviction. **a,** Comparison of promoter chromatin structure and chromatin accessibility by means of S5P CUTAC fragment size distribution over peaks (promoter and enhancer NDR spaces) in cells treated with DMSO (control) and Flavopiridol. Peaks were called with DMSO control. **b,** Heatmaps comparing nucleosomal (≥150 bp reads) and subnucleosomal (≤120 bp reads) protection by BAF (BRG1 CUT&RUN) and BAF-associated histones (BRG1 CUT&RUN.ChIP) in untreated (DMSO control) cells. Heatmaps were plotted relative to S5P CUTAC summits and sorted by decreasing accessibility (CUTAC signal). CUT&RUN.ChIP heatmaps show enrichment over IgG isotype control (for ChIP). **c, d,** Enrichment of nucleosomal (≥150 bp, solid lines) and subnucleosomal (≤ 120 bp, broken lines) reads from BRG1 CUT&RUN and CUT&RUN.ChIP experiments, relative to S5P CUTAC summits, in DMSO (c) and Flavopiridol treated (d) cells. **e,** Flavopiridol treatment causes eviction of partially unwrapped nucleosomes through enrichment of BAF at promoters and enhancers, leading to NDR persistence. Datasets are representative of at least two biological replicates.

We previously introduced the CUT&RUN.ChIP technique to demonstrate that the yeast counterpart of BAF, the RSC complex, binds partially unwrapped nucleosomes^5^. CUT&RUN.ChIP used MNase-digested chromatin released by RSC CUT&RUN as input for subsequent chromatin immunoprecipitation (ChIP) of histone epitopes, which revealed RSC-associated nucleosome substructures^5,38^. We used this method to ask whether BAF is similarly associated with unwrapped nucleosomal intermediates in mESCs. By analyzing nucleosomal (≥150 bp) and subnucleosomal (≤120 bp) sized DNA fragments protected from MNase digestion in BRG1 CUT&RUN.ChIP, we determined that subnucleosomal particles protecting <120 bp of DNA over the S5P CUTAC peaks are BAF-associated partially unwrapped nucleosomal intermediates (Fig. 2b,c). BAF-associated partially unwrapped nucleosomes were also observed immediately upstream of TSSs within gene promoters, consistent with their presence within promoter NDRs (Extended Data Fig. 3c). In contrast, nucleosomes flanking the CUTAC peaks and promoter NDRs were fully wrapped and protected >150 bp of DNA from MNase digestion (compare solid versus broken lines). BAF-associated partially unwrapped nucleosomes were enriched for histone H3K4me1 and H3K4me3 as well as the repressive H3K27me3 catalyzed by the Polycomb Repressive complex 2 (PRC2). BAF binding of H3K27me3 nucleosomes is consistent with its role in evicting H3K27me3 and PRC2 to activate transcription^39^.

Strikingly, treating mESCs with Flavopiridol resulted in a dramatic depletion of the partially unwrapped nucleosomal intermediates over S5P CUTAC peaks and promoter NDRs, while the flanking fully wrapped nucleosomes were retained (Fig. 2d, Extended Data Fig. 3d). Analysis of RNAPII-S5P CUTAC fragment sizes upon Flavopiridol treatment for eight hours showed a shift towards shorter fragments, implying enhanced DNA accessibility and indicating that partially unwrapped nucleosomes are evicted, although total S5P CUTAC signals remained the same (Fig. 2a, Extended Data Fig. 3a,b). Taken together, BRG1 CUT&RUN.ChIP and RNAPII-S5P CUTAC before and after Flavopiridol treatment suggest that enrichment of RNAPII pausing and subsequently elevated BAF occupancy leads to increased nucleosome eviction from gene promoters and distal accessible sites to establish steady-state NDRs (Fig. 2e). This provides a mechanistic basis for chromatin accessibility and shows how transcription and RNAPII drives the maintenance of accessible promoter and enhancer DNA by facilitating an ATP-dependent chromatin remodeler.

### RNAPII and BAF dynamically probe Polycomb-repressed chromatin

In multicellular eukaryotes, genes that are marked for repression are packaged into constitutive or facultative heterochromatin, which are nucleosome-dense regions and contain histone H3 with trimethylation of lysine 9 (H3K9me3) or lysine 27 (H3K27me3), respectively. In ESCs, H3K27me3 enriched genes are “bivalent” as they also contain the active H3K4me3 modification at promoters^40^ (Fig. 3a,b). These developmentally repressed genes are either transcriptionally inactive or transcribed at a very low level, and are considered to be “poised” for activation upon receiving differentiation signals. To compare transcriptionally active and bivalent chromatin regions genome-wide, we grouped mESC genes as RNAPII enriched (active) and H3K27me3 enriched (bivalent) (Extended Data Fig. 4a, Fig. 3b). CUT&Tag showed very low occupancy of RNAPII-S5P and BAF (BRG1) at bivalent promoters, but enrichment of H3K27me3 over the promoter and gene-body regions (Fig. 3a,b). Consistent with previous work, RNAPII-S5P CUTAC showed much reduced DNA accessibility at bivalent promoters compared to active promoters (Fig. 3b)^41^.

**Fig. 3.**
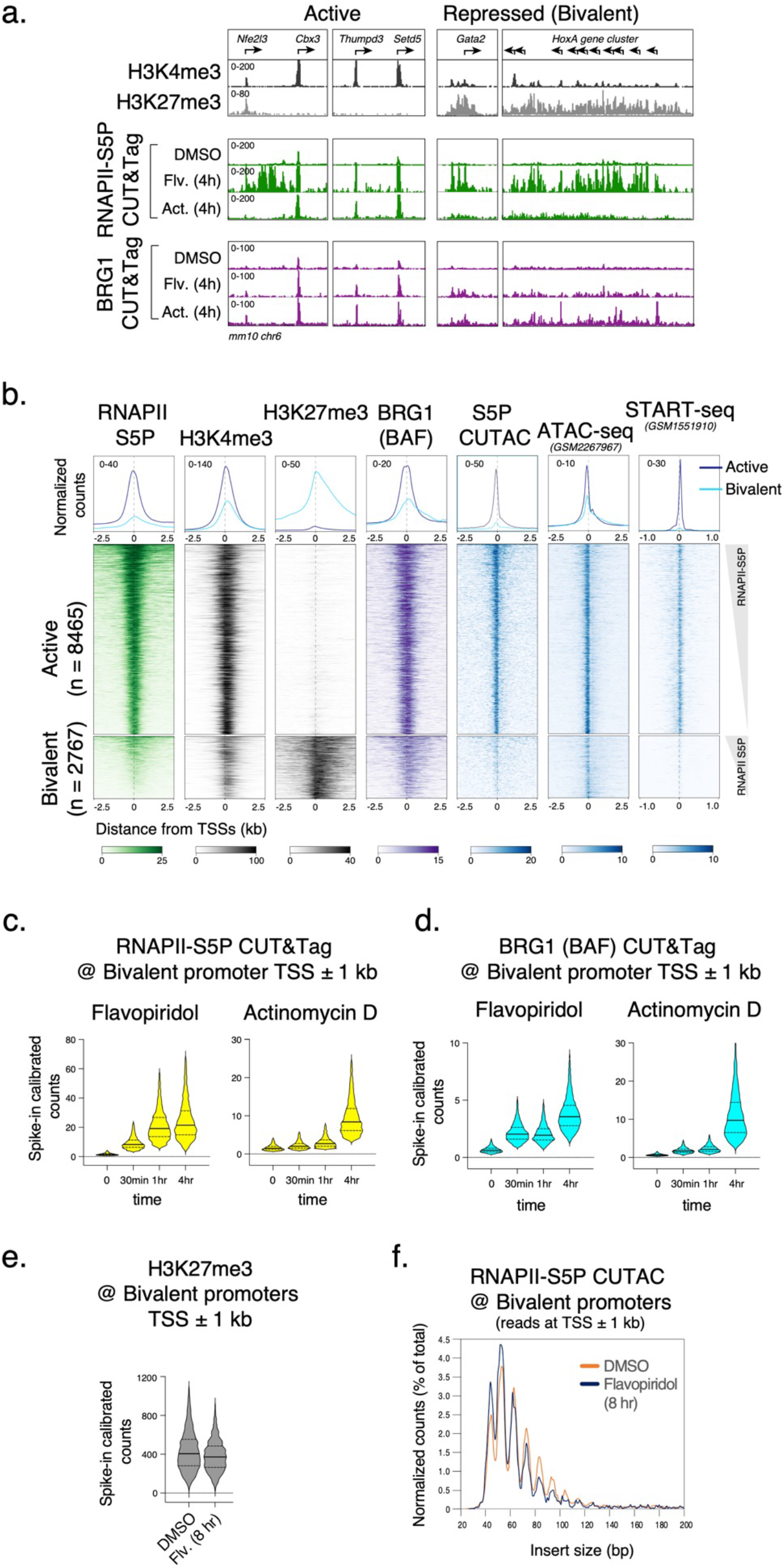
Transcription inhibition show RNAPII and BAF occupancy at Polycomb-repressed gene promoters. **a,** Representative genomic tracks comparing enrichment of histone modifications, RNAPII-S5P, and BRG1 by CUT&Tag at transcriptionally active and Polycomb-repressed bivalent genes, and changes in RNAPII-S5P and BRG1 occupancy upon Flavopiridol (Flv.) and Actinomycin D (Act.) treatment. RNApII-S5P and BRG1 CUT&Tag read counts were spike-in calibrated. **b,** Heatmaps (bottom) and average plots (top) comparing histone modifications (CUT&Tag), chromatin structure (RNAPII-S5P CUTAC and ATAC-seq), and transcriptional activity (START-seq) at RNAPII-enriched (active) and H3K27me3-enriched (bivalent) promoters. Promoters were grouped based on K-means clustering of RNAPII-S5P and H3K27me3 CUT&Tag reads mapping to a 5 kb window around the TSSs of RefSeq-annotated mESC genes, see Extended Data Fig. 4a. **c, d,** Violin plots of spike-in calibrated CUT&Tag signal distribution comparing RNAPII-S5P and BRG1 occupancy over bivalent promoter TSSs ± 1 kb at time points after drug treatments. Median value (solid line), upper- and lower quartiles (broken lines) and outliers were calculated using the Tukey method. **e,** Violin plot comparing spike-in calibrated H3K27me3 CUT&Tag at bivalent promoter TSSs ± 1 kb in cells treated with DMSO and Flavopiridol. **f,** S5P CUTAC fragment size distribution to compare chromatin accessibility at bivalent promoter TSSs ± 1 kb in cells treated with DMSO and Flavopiridol. Datasets are representative of at least two biological replicates.

Interestingly, treating cells with Flavopiridol or Actinomycin D over a time course resulted in a gradual buildup of RNAPII-S5P and concomitantly BAF at the bivalent (H3K27me3-high) promoters, mirroring the effects observed at active (RNAPII-high) promoters (Fig. 3a,c,d, Extended Data Fig. 4b,c). These include classic examples of Polycomb repressed genes such as the *Hox* gene clusters. Since BAF is known to oppose Polycomb repression^39^, we investigated whether increased BAF occupancy is associated with any effect on H3K27me3 and chromatin accessibility at bivalent promoters. CUT&Tag mapping showed a slight decrease in PRC2-catalyzed H3K27me3 upon eight hours of Flavopiridol treatment (Fig. 3e, Extended Data Fig. 4d). RNAPII-S5P CUTAC also showed a slight difference in fragment size distribution (Fig. 3f), suggesting that increased RNAPII-S5P and BAF occupancy is not sufficient for persistent chromatin remodeling and NDR establishment at bivalent promoters.

These data also show that RNAPII and BAF are not excluded from bivalent genes in mESCs. Rather, it is likely that that RNAPII and BAF continuously probe Polycomb-repressed chromatin and transiently initiate transcription. This is not unprecedented, as low levels of transcripts from Polycomb-repressed chromatin are detectable^42^, and Polycomb mediated repression was shown to reduce RNAPII burst frequency^43^. In contrast to the Polycomb-repressed facultative heterochromatin regions, RNAPII and BAF appear to be excluded from constitutive heterochromatin characterized by the H3K9me3 histone modification, where we do not observe occupancy even upon the drug treatments, in comparison to active and bivalent chromatin regions (Extended Data Fig. 4e).

### DNA-sequence-specific TFs capture transient site exposure to facilitate nucleosome eviction

If BAF and RNAPII probe both transcriptionally active and Polycomb-repressed chromatin, what determines their specificity for persistent chromatin remodeling to establish NDRs? In mESCs, these regions have differential binding of certain DNA-sequence-specific TFs such as NANOG, SOX2, OCT4 (POU5F1) and KLF4. These TFs are master regulators of ESC self-renewal and pluripotency and occupy almost all transcriptionally active promoters and enhancers (Fig. 4a, Extended Data Fig. 5a). Therefore, a parsimonious explanation is that chromatin binding of these pluripotency TFs drives nucleosome eviction and NDR establishment. Mechanistically, pluripotency TFs can bind nucleosomes *in vitro*^44^ and maintain chromatin accessibility *in vivo*^45^. However, ATP-dependent remodeling by BAF is in turn required to establish chromatin accessibility for pluripotency TF binding for ESC self-renewal^9,46–48^, during reprogramming^49^, and in developing blastocysts^50^. Therefore, how BAF and pluripotency TFs function to establish ESC-specific chromatin structure and pluripotency transcription network has remained a conundrum.

**Fig. 4.**
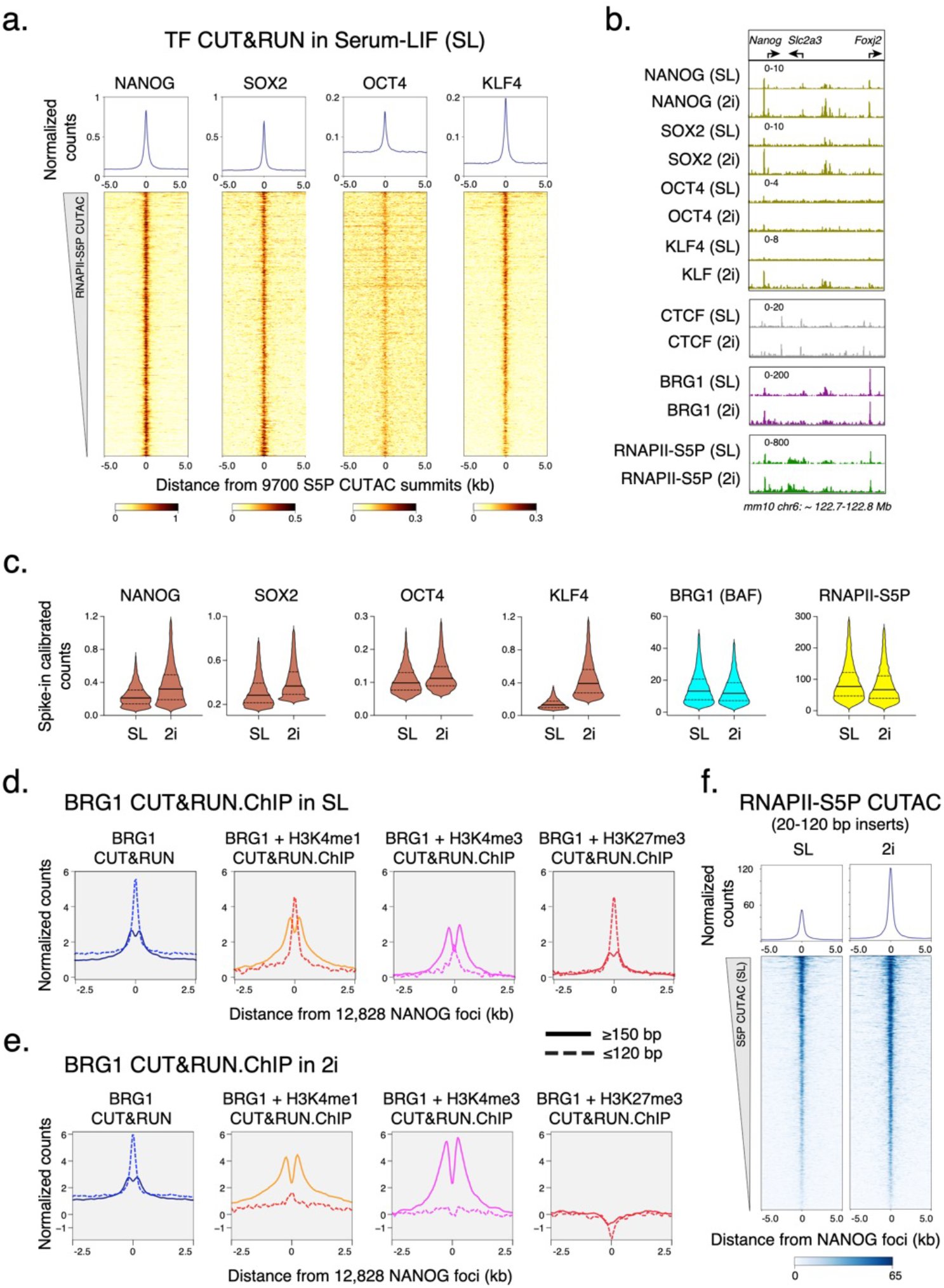
Pluripotency TF binding drives NDR establishment in mESCs. **a,** Heatmaps (bottom) and average plots (top) of pluripotency TF CUT&RUN reads relative to RNAPII-S5P CUTAC summits, showing TF binding at sites of DNA accessibility. Heatmaps were sorted by decreasing accessibility (CUTAC signal). **b,** Representative genomic tracks comparing occupancy of TFs (CUT&RUN), BRG1 (CUT&Tag), and RNAPII-S5P (CUT&Tag) in SL and 2i conditions. All datasets were spike-in calibrated. **c,** Violin plots of spike-in calibrated CUT&RUN (TF) and CUT&Tag (BRG1 and RNAPII-S5P) signal distribution comparing factor occupancy over RNAPII-S5P CUTAC peaks in 2i versus SL. Median value (solid line), upper- and lower quartiles (broken lines) and outliers were calculated using the Tukey method. **d, e,** Enrichment of nucleosomal (≥150 bp, solid lines) and subnucleosomal (≤ 120 bp, broken lines) reads from BRG1 CUT&RUN and CUT&RUN.ChIP experiments, relative to NANOG foci (smallest fragment within primary peaks called in SL condition), in SL (d) and 2i (e). **f,** Heatmaps (bottom) and average plots (top) of RNAPII S5P CUTAC (20-120 bp reads only) relative to NANOG foci, comparing chromatin accessibility in 2i versus SL. Datasets are representative of at least two biological replicates.

We used CUT&RUN to map pluripotency TF binding in mESCs, and found tight correspondence between RNAPII-S5P CUTAC and these TFs (Fig. 4a,b). To investigate how pluripotency TFs regulate nucleosome dynamics and chromatin accessibility (NDR establishment), we used culture conditions to modulate TF protein concentration in mESCs. Culturing mESCs in media containing serum and Leukemia Inhibitory Factor (LIF) mimics a post-implantation embryonic stage (embryonic day 4.5) and ensures expression of the pluripotency TFs^51^. However, protein levels of NANOG and KLF4 are heterogeneous in individual cells cultured in this “naïve state” pluripotency condition (called serum-LIF or SL)^52,53^. Dual inhibition of the signaling kinases GSK3 and MEK together with LIF promotes a cellular state closer to the pluripotent pre-implantation epiblast (embryonic day 3.5)^54,55^. In this “ground state” pluripotency condition (called 2i), cells upregulate NANOG expression resulting in a higher and homogenous protein level in individual mESCs^56^. Immunofluorescent imaging confirmed strong upregulation of NANOG and KLF4 in 2i compared to SL (Extended Data Fig. 5b). Consistent with higher protein concentration in cells, CUT&RUN showed a more than three-fold increase in chromatin binding of KLF4, and a modest increase for NANOG and SOX2 (Fig. 4b,c, Extended Data Fig. 5c). In contrast, we did not observe significant changes in chromatin binding of OCT4 and of the ubiquitous TF insulator protein CTCF. Interestingly, CUT&Tag mapping for BRG1 showed that chromatin occupancy of BAF is comparable in SL and 2i (Fig. 4b,c, Extended Data Fig. 5c), which implies that pluripotency TFs do not recruit BAF to chromatin.

We showed by CUT&RUN.ChIP mapping of BRG1 and histone co-occupancy that BAF is associated with partially unwrapped nucleosomes at promoters and distal S5P CUTAC sites (Fig. 2b,c) which overlap with pluripotency TF binding (Fig. 4a). When plotted directly over the binding sites (peaks) of a pluripotency TF, *e.g*., NANOG, BRG1 CUT&RUN.ChIP showed similar partially unwrapped nucleosomal intermediates (Fig. 4d). These data suggest that nucleosome unwrapping by BAF facilitates pluripotency TF binding within gene regulatory elements in mESCs, as we have previously proposed for the yeast GRFs Abf1 and Reb1^5^. We next compared BRG1 CUT&RUN.ChIP in 2i (higher TF expression) vs SL (lower TF expression). We observed a striking loss of BAF-associated partially unwrapped nucleosomes over NANOG and S5P CUTAC peaks in 2i compared to SL (Fig. 4e), similar to what we observed upon treating cells with Flavopiridol in SL (Fig. 2d). Interestingly, S5P CUTAC showed a two-fold increase in chromatin accessibility in 2i compared to SL (Fig. 4f), despite comparable RNAPII-S5P occupancy in the two conditions (Fig. 4b,c). We conclude that pluripotency TFs capture transient site exposure due to nucleosome unwrapping by BAF to further destabilize and evict nucleosomes, and increased TF abundance drives this process towards more stable NDR formation, consistent with the robustness of the 2i culture condition for ESC pluripotency maintenance.

## Discussion

We have demonstrated that RNAPII promoter-proximal pausing strongly facilitates BAF chromatin binding and nucleosome remodeling. We also showed that RNAPII and BAF dynamically probe both transcriptionally active euchromatin and Polycomb-repressed facultative heterochromatin in mESCs. We found that nucleosome eviction by BAF to establish NDRs additionally requires pluripotency TF binding. Our study suggests that the mechanism establishing and maintaining steady-state chromatin accessibility patterns involves a functional synergy between RNAPII promoter-proximal pausing, BAF and DNA-sequence-specific TFs (Fig. 5). In our model, TF-chromatin binding is the determinant step converting abortive BAF remodeling (discussed below) into productive nucleosome eviction and NDR establishment, thereby acting as a switch between transcriptionally repressive and transcriptionally active chromatin.

**Fig. 5.**
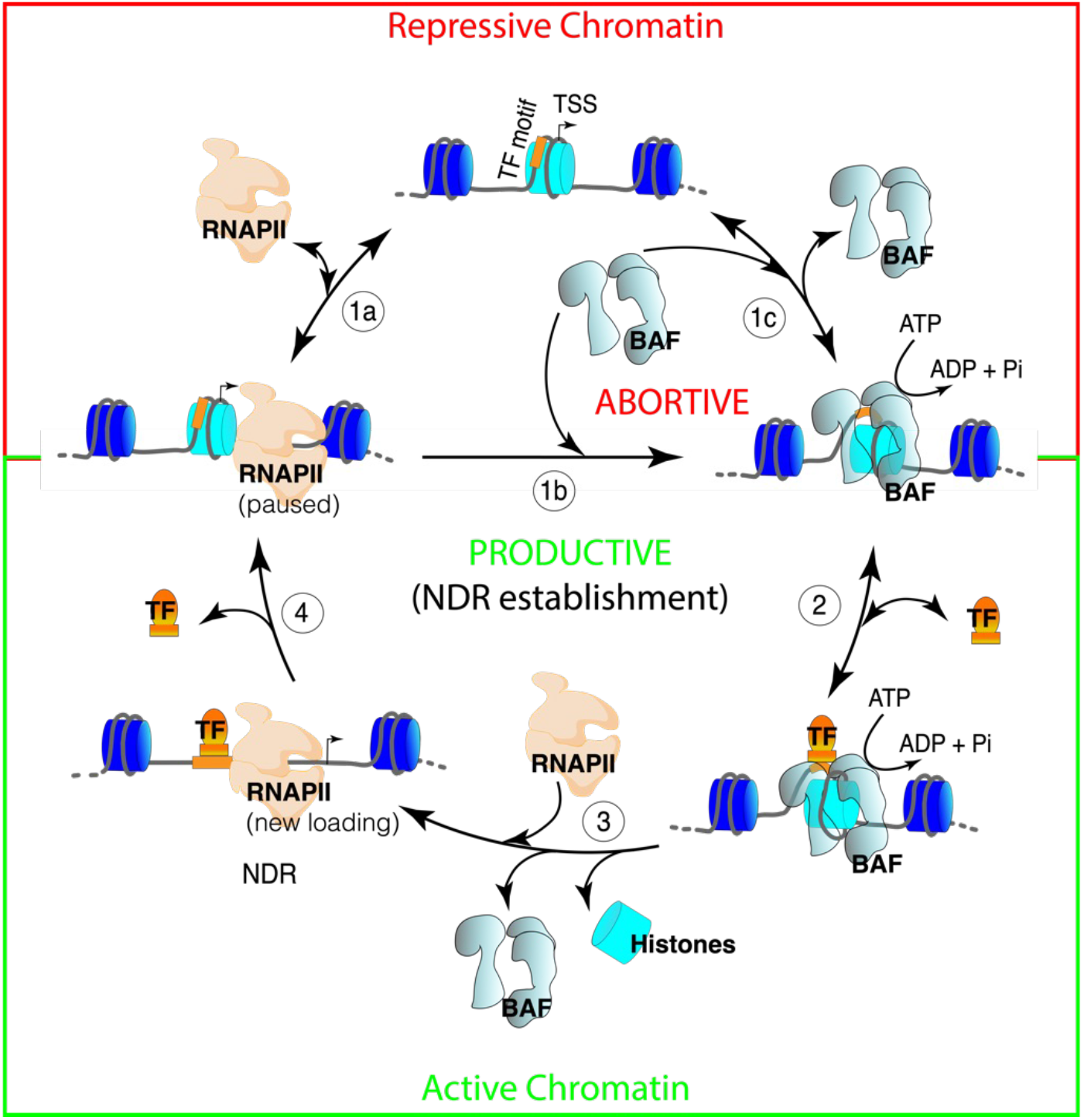
RNAPII, BAF and DNA-sequence-specific TFs synergize to dynamically generate NDRs. Model showing that RNAPII and BAF dynamically engage chromatin in an abortive manner (steps 1a-c) and require chromatin binding by DNA-sequence-specific TFs for productive chromatin remodeling and histone eviction to form an NDR (steps 2-4). Productive remodeling is essential to establish and maintain an active chromatin structure. Steady state promoter and enhancer chromatin structure is highly dynamic and constitutes a cycle of nucleosome deposition and eviction as well as factor binding and unbinding. In this cycle, RNAPII promoter proximal pausing facilitates BAF binding (step 1b). BAF translocates along nucleosomal DNA using the energy of ATP and partially unwraps nucleosomes occluding TF motifs, but BAF turns over rapidly (step 1c). TFs capture transient site exposure to bind to partially unwrapped nucleosomes (step 2), and synergize with BAF to evict histones and create an NDR (step 3), which is essential for subsequent loading of new RNAPII and transcription initiation. TFs also have seconds residence times on chromatin and turn over rapidly (step 4), such that a nucleosome can be deposited or moved into the NDR space, resuming the cycle. Increased expression of TFs (as in 2i compared to SL mESC culture condition) or higher TF DNA binding affinity drives steps 2 and 3 towards a more nucleosome-depleted state by making step 2 irreversible.

Positioned nucleosomes are refractory to RNAPII loading, which requires chromatin remodeling. However, chromatin undergoes major restructuring during DNA replication as nucleosomes are disassembled ahead of the replication fork and are replaced randomly on nascent DNA^57,58^. We hypothesize that RNAPII broadly scans chromatin for exposed DNA and utilizes the window of opportunity in the wake of DNA replication to bind to promoter and enhancer regions and initiate transcription (Fig. 5, step 1a). RNAPII promoter-proximal pausing would then facilitate BAF binding (step 1b) to clear away nucleosomes that encroach into the NDR space upstream of the paused RNAPII to ensure subsequent rounds of RNAPII loading. Consistent with our model, analysis of newly-replicated chromatin shows that transcription begins early, before chromatin maturation, and a delay in the maturation of repressive chromatin provides a temporal window for TF binding and chromatin activation^58,59^. BAF translocates on nucleosomal DNA using the energy of ATP hydrolysis and breaks histone-DNA contacts to partially unwrap a nucleosome. We hypothesize that BAF rapidly unbinds (step 1c) even before nucleosome eviction, which may involve multiple rapid cycles of BAF binding, nucleosomal DNA translocation, and unbinding events. Live-cell imaging of *Drosophila* Brahma and yeast RSC remodelers support our hypothesis of abortive BAF remodeling and show that chromatin binding of SWI/SNF remodelers is strikingly dynamic with less than 5 seconds average residence and turnover times, whereas histone turnover is more than two orders of magnitude slower^60,61^. Intriguingly, the fast dissociation kinetics of the remodelers is dependent on ATP-hydrolysis, suggesting that dissociation is a part of the remodeling mechanism. *In vitro* single-molecule measurements using physiological ATP concentrations have estimated that on average, nucleosomal DNA translocation rate of SWI/SNF remodelers is 12 bp/s^62^. This translocation speed combined with a short chromatin residence time agrees with multi-turnover remodeling for nucleosome eviction, which may require a major part of the 147 bp nucleosomal DNA to be disrupted, particularly at the histone H3-H4 interface. In a simplified two-component system (BAF and nucleosomes) where ATP is not limiting, the kinetics of nucleosome eviction would therefore be an outcome of a dynamic competition between nucleosome re-wrapping and BAF re-binding. Consistent with our model, chemically-induced proximity-mediated tethering of BAF to Polycomb-repressed promoters was shown to rapidly evict H3K27me3 and increase chromatin accessibility in an ATP-dependent manner^63^. We found that paused RNAPII facilitates BAF binding to drive nucleosome eviction. However, paused RNAPII is also distinctively dynamic and rapidly turns over in a seconds timescale (step 1a), primarily due to premature termination, as shown by live-cell imaging as well as genome-wide mapping and single-molecule footprinting experiments^35,64,65^. Taken together, the short residence times of BAF and paused RNAPII suggest the requirement for additional factors or mechanisms for productive nucleosome eviction and NDR establishment, without which BAF and RNAPII functions are abortive.

We found that BAF-associated partially unwrapped nucleosomes in mESCs are enriched at pluripotency TF binding sites within active gene promoters and enhancers, suggesting that nucleosomes occluding the TF-binding sites are remodeled by BAF. These dynamic nucleosomes are highly sensitive to MNase digestion in comparison to MNase-resistant nucleosomes genome-wide^66^. At their target loci, TFs could capture DNA sequence motifs transiently exposed due to BAF remodeling to further destabilize and evict nucleosomes, forming an NDR (steps 2-3)^2,67,68^. Indeed, we had previously shown that the yeast GRFs Abf1 and Reb1 bind to partially unwrapped nucleosomes that are targets of RSC remodeling^5^, as expected if increased TF concentration or TF DNA-binding affinity drives the nucleosome depleted state^67^. In agreement with this model, increased expression of mESC-specific pluripotency TFs NANOG and KLF4 in 2i compared to SL culture conditions resulted in enhanced nucleosome eviction over the TF-binding sites, showing how BAF and TFs synergize to establish locus-specific chromatin accessibility. TF binding may therefore provide an energetic advantage by reducing the ATP expense associated with abortive BAF remodeling. In contrast, Polycomb-repressed heterochromatin does not contain pluripotency TF binding motifs. Consequently, transient RNAPII and BAF activity is not sufficient for productive nucleosome remodeling and eviction to establish chromatin accessibility at Polycomb-repressed bivalent promoters. However, this dynamic between BAF and RNAPII does not occur in constitutive heterochromatin, which is maintained in a silent state by heterochromatin-associated proteins and DNA methylation.

ATP-dependent nucleosome remodeling by BAF is central to cell-type-specific transcriptional regulation in widely different tissues and across developmental processes^69,70^. Deregulation of BAF remodeling is linked with >20% of all human cancers and several neurodevelopmental disorders^25,71,72^. How BAF is targeted for precise spatiotemporal gene regulation, has remained an open question^73^. It is generally perceived that part of this problem is solved by cell-type-specific TFs recruiting BAF to specific target genomic loci. But whether and how TFs recruit BAF has remained unclear as a general mechanism. We found that increased pluripotency TF concentration to physiologically relevant levels and increased chromatin binding in mESCs has no effect on BAF occupancy, which argues against a recruitment-based mechanism. Rather, our model for coordination between BAF and TFs for nucleosome eviction is sufficient to explain the key roles of BAF in regulating locus- and cell-type-specific chromatin structure and transcription without the requirement for recruitment per se. Furthermore, our work broadly explains how perturbation of BAF dynamics and changes in cell-type-specific TF homeostasis in cancers may drive oncogenic gene expression programs. Although initial studies of BAF mutations in cancers identified BAF subunits as tumor suppressors, several recent findings suggest that BAF can also function as an oncogene. These include oncogenic fusion of the BAF subunit SS18 with an SSX protein in synovial sarcoma and interaction of BAF with EWS-FLI1 in Ewing sarcoma causing aberrant BAF complex targeting^74,75^, subunit overexpression^76^, and heterozygous missense mutations within the *Brg1* subunit gene in diverse cancers, several of which were characterized as gain-of-function for remodeling activity^77^. Furthermore, genetic dependency on *Brg1* observed in several cancers has led to the emergence of pharmacological inhibition of the BRG1 ATPase activity as a novel therapeutic strategy^78–80^. Our study explains how BAF can synergize with cell-type-specific chromatin regulators and TFs, shedding light into how BAF functions may be hijacked in cancers.

## Methods

### Cell culture

AB2.2 (ATCC SCRC-1023; Male; strain: 129S5/SvEvBrd) mouse embryonic stem cells were used in all experiments. Cells were thawed and initially cultured in “2i” media, consisting of Knockout Dulbecco’s modified Eagle’s medium (Gibco cat. No. 10829018), supplemented with 15% ES-qualified fetal calf serum (Gibco cat. No. 16141079), 2 mM L-glutamine (Sigma-Aldrich cat. No. G7513), 0.1 mM MEM non-essential amino acids (Gibco cat. No. 11-140-050), 0.1 mM β-mercaptoethanol (Gibco cat. No. 21985023), 1000 U/mL leukemia inhibitory factor (LIF) (MilliporeSigma cat. No. ESG1107), 3 μM GSK 3β inhibitor CHIR99021 (Sigma-Aldrich cat. No. SML1046), and 1 μM MEK inhibitor PD0325901 (Sigma-Aldrich cat. No. PZ0162). “Serum-LIF” or “SL” media contained all components except the GSK 3β and MEK inhibitors. Cells were maintained on 6-well plates or cell culture flasks coated with Attachment Factor Protein (Gibco cat. No. S006100) at 37°C with 5% CO2, passaged every 48-72 h with daily media changes. Cells were cultured for at least three passages before transferring to serum-LIF (SL) media and cultured for at least three more passages before experiments. Cells were released with Accutase (STEMCELL Technologies cat. No. 07922) for harvesting, washed with sterile phosphate buffered saline (PBS), resuspended in media supplemented with 10% dimethyl sulfoxide (DMSO) (Sigma-Aldrich cat. No. 41640), and slow-frozen at −80°C in isopropanol freezing chambers. All experiments were performed with cells harvested and frozen after six to eight total passages. Cultures were periodically tested for Mycoplasma and karyotyped to detect any chromosomal abnormalities. For inhibitor treatments, media was supplemented with 10 μM Triptolide (Selleckchem cat. No. S3604), 1 μM Flavopiridol hydrochloride hydrate (Sigma-Aldrich cat. No. FL3055), 5 ug/mL Actinomycin D (Sigma-Aldrich cat. No. A9415), or 1:1000 v/v DMSO, plates/flasks were transferred to ice at time points, cells were washed twice with ice-cold PBS and promptly harvested.

### CUT&Tag

CUT&Tag was performed using frozen cells as described previously^28^, with some modifications. Also see https://www.protocols.io/view/bench-top-cut-amp-tag-kqdg34qdpl25/v3 for a step-by-step protocol. For each CUT&Tag experiment, 0.2 x 10^6^ cells were bound to 10 μL of Bio-Mag Plus Concanavalin A coated magnetic beads (Bangs laboratories, cat. No. BP531), equilibrated with binding buffer (20 mM K-HEPES pH7.9, 10 mM KCl, and 1 mM each CaCl_2_ and MnCl_2_). Beads (with bound cells) were magnetized, supernatant removed, washed once with 400 μL, and resuspended in 200 μL of Wash buffer (20 mM Na-HEPES pH 7.5, 150 mM NaCl, 0.5 mM spermidine, and EDTA-free protease inhibitor) supplemented with 2 mM EDTA and 0.05% Digitonin (MilliporeSigma cat. No. 3004105GM). Primary antibodies were mixed at optimum dilutions (see below) and incubated overnight at 4°C on a rotating platform. Beads were washed once with 400 μL Dig-Wash (Wash buffer supplemented with 0.05% Digitonin), resuspended in 200 μL Dig-Wash with a secondary antibody (see below), and incubated for 30 min to 1 hour at room temperature (RT) on a rotating platform. Beads were washed twice with 400 μL Dig-Wash and resuspended in 200 μL Dig-Med buffer (Dig-Wash buffer, except containing 300 mM NaCl) with 1:200 dilution (~0.04 μM) of lab-made proteinA-Tn5 transposase fusion protein (pA-Tn5) pre-loaded with double-stranded adapters with 19-mer mosaic ends and containing carry over *E. coli* DNA, useful for spike-in calibration^28^. pA-Tn5 incubations were performed on a rotating platform for 1 hour at RT. Beads were washed three times with 400 μL Dig-Med to remove unbound pA-Tn5 and resuspended in 300 μL Tagmentation buffer (Dig-Med supplemented with 10 mM MgCl_2_). Tagmentation reactions were performed by incubating samples at 37°C on a rotating platform for 1 hour. Tagmentation reactions were stopped with 10 μL of 0.5 mM EDTA, 3.1 μL of 10% SDS (1% final), and 2 μL of 20 mg/mL Proteinase K (Invitrogen cat. No. 25530049) and incubated in a 50°C water bath for 1 hour or 37°C overnight with rotation. DNA was extracted using phenol-chloroform extraction method and precipitated using chilled 75% ethanol. DNA pellets were dissolved in 30 μL 0.1X TE (1 mM Tris pH 8, 0.1 mM EDTA) supplemented with 1:400 dilution of 10 mg/mL RNase A (Thermo Scientific cat. no. EN0531) and incubated in 37°C water bath for 15 min. Libraries were amplified using NEBNext HiFi 2X PCR Master mix (NEB cat. no. M0541) with 13 rounds of amplification as described previously^28^. Sequencing libraries were purified using a 1.3X ratio of HighPrep PCR Cleanup beads (MagBio genomics cat. no. AC-60500) as per manufacturer’s instructions and eluted in 0.1X TE. Library quality and concentration were evaluated using Agilent Tapestation D1000 capillary gel analysis.

### RNAPII-S5P CUTAC

CUTAC using RNAPII-S5P for accessible site mapping was performed as described in the step-by-step protocol https://www.protocols.io/view/cut-amp-tag-direct-with-cutac-x54v9mkmzg3e/v3?step=119. Briefly, nuclei were prepared as previously described^28^ and lightly cross-linked (0.1% formaldehyde 2 min), then washed and resuspended in Wash buffer (10mM HEPES pH 7.5, 150 mM NaCl, 2mM spermidine, and Roche complete EDTA-free protease inhibitor). CUTAC was performed with 0.05 x 10^6^ nuclei by mixing with 5 μL Concanavalin A magnetic beads. Primary antibody against RNAPII-S5P (Cell Signaling Technology cat. no. 13523) was added in 1:50 dilution in Wash buffer supplemented with 0.1% bovine serum albumin (BSA) and incubated overnight at 4 °C. Beads were magnetized, supernatant was removed, and beads were resuspended in Wash buffer containing 1:100 guinea pig anti-rabbit secondary antibody (Antibodies Online cat. no. ABIN101961) and incubated 0.5–1 hour at RT. Beads were magnetized and washed (on the magnet) once with Wash buffer, resuspended in pAG-Tn5 pre-loaded with mosaic-end adapters (EpiCypher cat. no. 15-1117, 1:20 dilution) in 300-Wash buffer (Wash buffer except containing 300 mM NaCl) and incubated 1 hour at RT. Beads were washed (on the magnet) three times in 300-Wash, then incubated at 37 °C for 20 min in 50 μL CUTAC-hex tagmentation solution (5 mM MgCl_2_, 10 mM TAPS, 10% 1,6-hexanediol). Bead suspensions were chilled on ice, magnetized, supernatant was removed, beads were washed with 10 mM TAPS pH 8.5, 0.2 mM EDTA, and resuspended in 5 μL 0.1% SDS, 10 mM TAPS pH 8.5. Beads were incubated at 58 °C in a thermocycler with heated lid for 1 hour, followed by addition of 15 μL 0.67% Triton X-100 to neutralize the SDS. Libraries were amplified using NEBNext HiFi 2X PCR Master mix with gap-filling and 12-cycle PCR: 58 °C 5 min, 72 °C 5 min, 98 °C 30 sec, 12 cycles of (98 °C 10 sec denaturation and 60 °C 10 sec annealing/extension), 72 °C 1 min, and 8 °C hold. Sequencing libraries were purified with 1.3X ratio of HighPrep PCR Cleanup beads as per manufacturer’s instructions and eluted in 0.1X TE. Library quality and concentration were evaluated using Agilent Tapestation D1000 capillary gel analysis.

### CUT&RUN and CUT&RUN.ChIP

BRG1 CUT&RUN.ChIP was performed as described previously^5,38^ with some modifications. For CUT&RUN, 2.5 x 10^6^ cells were bound to 50 μL of Concanavalin A coated magnetic beads. Primary and secondary antibody incubation and washes were performed as described above for CUT&Tag, but in 1 mL of the buffers using 1.5 mL low-binding flip-cap tubes. Incubations were done at 4°C, and ice-cold buffers were used in every step. After secondary antibody incubation and washes, bead-bound cells were resuspended in ice-cold Dig-Wash with lab-made proteinA-micrococcal nuclease fusion protein (pA-MN, 360 μg/ml, 1:400 dilution) and incubated 1 hour at 4°C with rotation. The beads were washed three times in ice-cold Dig-Wash, resuspended in 0.5 mL ice-cold Dig-Wash, and equilibrated to 0°C. CaCl_2_ was quickly mixed to a final concentration of 2 mM, incubated on ice for 5 min for MNase digestion, and reactions were stopped with 0.5 mL of 2XSTOP buffer (150 mM NaCl, 20 mM EDTA, 4 mM EGTA, and 50 μg/mL RNase A) supplemented with BRG1 peptide (Abcam cat. no. ab241115) to a final concentration of 10 μg/mL. Samples were incubated at 37°C for 20 min and centrifuged for 5 min at 16,000 x g and 4°C. The supernatant was removed on a magnet stand and divided into five 200 μL aliquots for ChIP. One aliquot was saved (at 4°C) as the input. To the ChIP samples, respective antibodies (IgG and histone PTMs, see below) were added and incubated at 4°C overnight. Protein A Dynabeads (Invitrogen cat. no. 10002D) were equilibrated in Wash buffer supplemented with 0.05% Tween-20, and 20 μl of beads were added to each ChIP sample (except the input). Samples were incubated at 4°C for 30 min and washed once with Wash buffer+Tween-20. The ChIP samples were brought up with DNA-extraction buffer (150 mM NaCl 10 mM EDTA, 2 mM EGTA, 0.1% SDS, and 0.2 mg/ml Proteinase K). SDS (0.1%) and Proteinase K (0.2 mg/ ml) were added separately to the input samples. Samples were incubated at 50°C for 1 hour. DNA was extracted using phenol-chloroform extraction method and 40 μg of glycogen (Roche cat. no. 10901393001) was mixed with the aqueous phase. DNA was precipitated using chilled 75% ethanol and dissolved in 0.1X TE.

For each TF CUT&RUN, 0.5 x 10^6^ cells were bound to 10 μL of Concanavalin A coated magnetic beads. Incubations and washes were done as for BRG1 CUT&RUN, except that primary and secondary antibody incubations and pA-MN binding were performed in 200 μL volumes, the MNase digestion reaction was done in 150 μL with incubation on ice for 30 min, and the reaction was stopped using 150 μL of 2XSTOP buffer without any peptide. Samples were incubated at 37°C for 20 min and centrifuged for 5 min at 16,000 x g and 4°C. The supernatant containing released chromatin particles was removed on a magnet stand and SDS (0.1%) and Proteinase K (0.2 mg/mL) were added. Samples were incubated at 50°C for 1 hour and used directly for DNA extraction using the phenol-chloroform extraction method described above. Libraries were prepared for Illumina sequencing with Tru-Seq adapters as described^5,38^. Briefly, libraries were constructed without size-selection, following the KAPA DNA polymerase library preparation kit protocol (https://www.kapabiosystems.com/product-applications/products/next-generation-sequencing-2/dna-library-preparation/kapa-hyper-prep-kits/), optimized to favor exponential amplification of <1000 bp fragments over linear amplification of large DNA fragments. Sequencing libraries were then purified using 1.2X ratio of HighPrep PCR Cleanup System. Library concentrations were quantified using the D1000 TapeStation system (Agilent).

### Sequencing, data processing, data analysis, and data visualization

Libraries were sequenced for 25 cycles in 25 bp paired-end mode on the Illumina HiSeq 2500 or 50 bp paired-end on NextSeq 2000 at the Fred Hutchinson Cancer Center Genomics Shared Resource, and data were analyzed as described (https://www.protocols.io/view/cut-amp-tag-data-processing-and-analysis-tutorial-e6nvw93x7gmk/v1). Briefly, adapters were clipped and paired-end Mus musculus reads were mapped to UCSC mm10 using Bowtie2^81^ with parameters: --very-sensitive-local --soft-clipped-unmapped-tlen --dovetail --no-mixed --no-discordant -q --phred33 -I 10 -X 1000 (for CUT&Tag) or --end-to-end --very-sensitive --no-mixed --no-discordant -q --phred33 -I 10 -X 700 (for CUT&RUN.ChIP) . Spike-in *E. coli* reads in CUT&TAG experiments were mapped to Ensembl masked R64-1-1 with parameters: --end-to-end --very-sensitive --no-overlap --nodovetail --no-unal --no-mixed --no-discordant -q --phred33 -I 10 -X 700. Continuous-valued data tracks (bedGraph and bigWig) were generated using genomecov in bedtools v2.30.0 (-bg option) and normalized as fraction of total counts (for CUTAC and CUT&RUN.ChIP) or calibrated using total number of spike-in reads (for CUT&Tag)^28^. Genomic tracks were displayed using Integrated Genome Browser. RNAPII-S5P CUTAC H3K9me3 CUT&Tag and TF CUT&RUN peaks were called by SEACR (v.1.3) using the norm and relaxed settings^82^; 20-120 bp fragments were used to call TF peaks. Profile plots, heatmaps, and correlation matrices were generated using deepTools v3.5.1^83^. Scores were averaged over 50 bp non-overlapping bins with respect to reference points and plotted as the mean. Violin plots were generated with GraphPad Prism 9. Scores were computed using deepTools v3.5.1, and extreme outliers were identified using the ROUT method (Q = 0.2%) and removed.

### Immunofluorescence

Cells were cultured as described above and immunofluorescence staining was conducted in-well using 12-well plates at RT. After culturing, cells were washed once with 1 mL PBS with gentle rocking for 5 minutes, then incubated with 4% paraformaldehyde in 1 mL PBS for 15 minutes with gentle rocking. Wells were rinsed once with PBS, then washed twice with 1 mL PBS supplemented with 0.1% Triton X-100 (PBST) for 5 minutes each with gentle rocking. Wells were then incubated with 0.5 mL PBST containing primary antibody in optimum dilution (see below) and 1% BSA, overnight at 4°C with gentle rocking. Wells were rinsed once, then washed twice for 5 minutes each with 1 mL PBST. Wells were then incubated with 0.5 mL PBST containing fluorophore-conjugated secondary antibody (see below) for 1 hour at RT with gentle rocking. Wells were rinsed once and washed twice for 5 min each with 1 mL PBST, then incubated with 0.5 mL PBST with 1:50,000 4’,6-diamidino-2-phenylindole (DAPI) for 20 min at RT with gentle rocking for nucleic acid staining. Wells were rinsed once and washed three times for 5 min each with 1 mL PBST, then imaged in 0.5 mL PBS on an EVOS FL Auto 2 Cell Imaging System (Invitrogen) with 10x magnification.

### Antibodies

RNAPII-S5P: rabbit monoclonal (Cell Signaling Technology cat. no. 13523), 1:50; RNAPII-S2P: rabbit monoclonal (Cell Signaling Technology cat. no. 13499), 1:100; RPB3: rabbit polyclonal (Bethyl Laboratories cat. no. A303-771A, lot no. A303-771A2), 1:100; BRG1: rabbit monoclonal (abcam cat. no. ab110641), 1:100, for CUT&Tag and CUT&RUN.ChIP, and rabbit polyclonal (Invitrogen cat. no. 720129, lot. no. 2068859), 1:250, for immunofluorescence; H3K4me1: rabbit polyclonal (Abcam cat. no. ab8895, lot no. GR3283237), 1:100; H3K4me3: rabbit polyclonal (Active Motif cat. no. 39915, lot. no. 24118008), 1:100, for CUT&Tag, and rabbit monoclonal (EpiCypher cat. no. 13-0028), 1:100, for CUT&RUN.ChIP; H3K27me3: rabbit monoclonal (Cell Signaling Technology cat. no. 9733), 1:100; H3K9me3: rabbit monoclonal (Abcam cat. no. ab176916), 1:100; guinea pig anti-rabbit secondary: Antibodies Online cat. no. ABIN101961, 1:100; isotype control (IgG) for CUT&RUN. ChIP: rabbit monoclonal (abcam cat. no. ab172730), 1:100; NANOG: rabbit polyclonal (Bethyl Laboratories cat. no. A300-397A, lot no. 3), 1:100; KLF4: goat polyclonal (R&D Systems cat. no. AF3158, lot. no. WRR0719011), 1:100 for CUT&RUN, 1:50 for immunofluorescence; OCT4: rabbit monoclonal (abcam cat. no. ab181557), 1:100; SOX2: rabbit monoclonal (Abcam cat. no. ab192494), 1:100; CTCF: rabbit monoclonal (Abcam cat. no. ab128873), 1:100; rabbit anti-goat secondary: abcam cat. no. ab6697, 1:100; goat anti-rabbit-Cy5 secondary: Jackson ImmunoResearch Cat. no. 111-175-144, 1:500; donkey anti-goat-rhodamine red secondary: Jackson ImmunoResearch cat. no. 705-295-147, 1:250.

## Data availability

All primary sequencing data have been deposited as paired-end fastq files and all mapped data have been deposited as bigWig files in the Gene Expression Omnibus under the accession number GSEXXXXXXX.

Public datasets used: ATAC-seq: GSM2267967^84^; START RNA-seq: GSM1551910^30^; MNase seq: GSE117767^41^; mESC Enhancer annotation: Whyte et al., 2013^85^

## Acknowledgements

We thank Kami Ahmad and Toshi Tsukiyama for critical readings of the manuscript, Jorja Henikoff for help with processing of sequencing data, and David Scalzo and and Xiaozhong Wang (Northwestern University) for guidance on mESC culturing. This research was supported by NIH K99 GM138920 (S.B.), the Howard Hughes Medical Institute (S.H.), and NIH P30CA015704 (Fred Hutch Shared Resources)

## Author Information

### Authors and Affiliations

Basic Sciences Division, Fred Hutchinson Cancer Research Center, Seattle, WA, USA: Sandipan Brahma, Steven Henikoff

Howard Hughes Medical Institute, Chevy Chase, MD, USA: Steven Henikoff

### Contributions

Conceptualization, S.B.; Investigation, S.B.; Writing – original draft, S.B.; Writing – Review & Editing, S.B., and S.H.; Funding Acquisition, S.H., and S.B.; Resources, S.H. Both authors approved the final manuscript.

### Corresponding authors

Correspondence to Sandipan Brahma, Steven Henikoff

## Extended Data Figures

**Extended Data Fig. 1.**
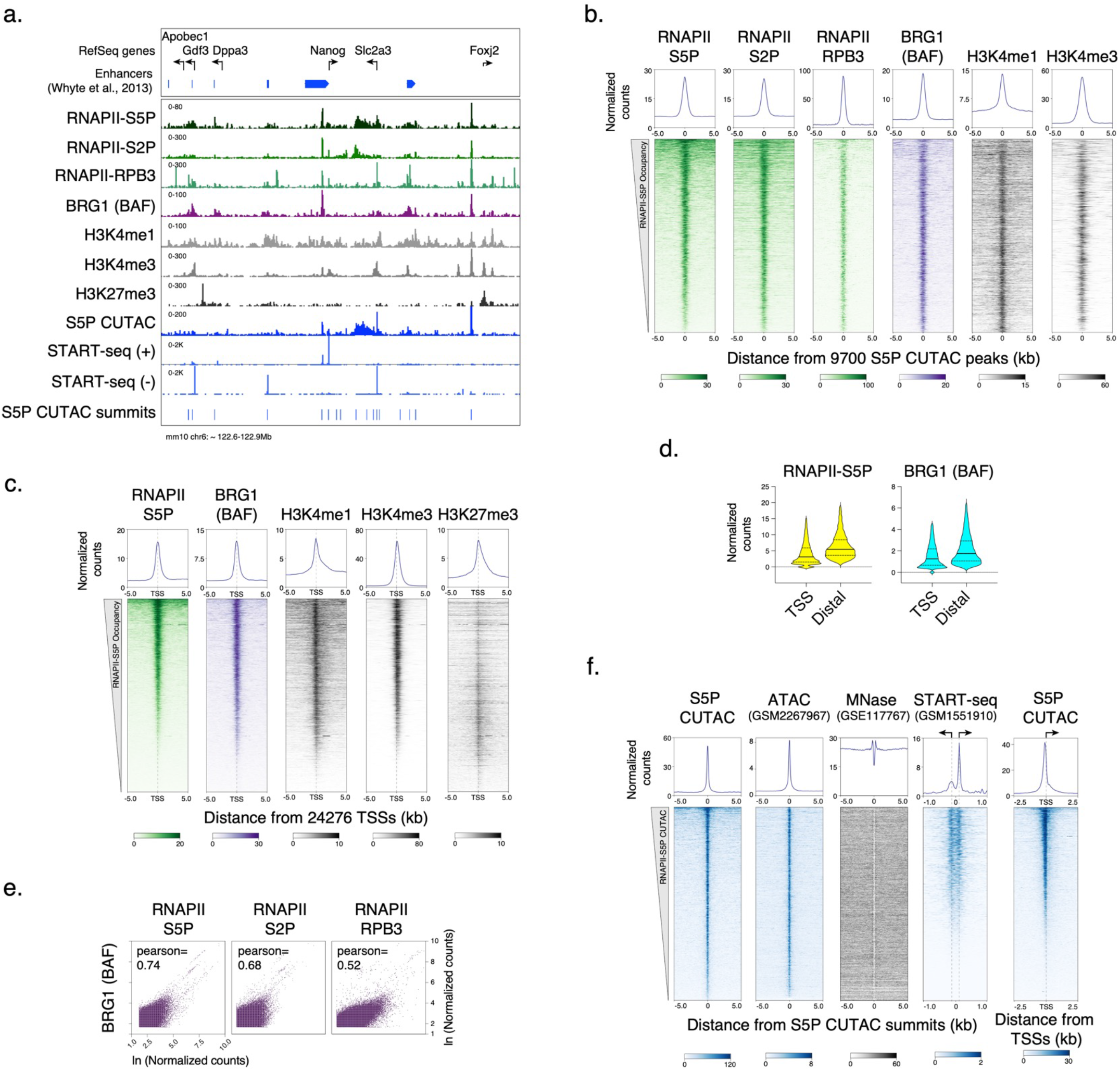
CUT&Tag of chromatin epitopes and RNAPII-S5P CUTAC in mESCs. **a,** Representative genomic tracks showing RNAPII, BRG1, histone PTM occupancy by CUT&Tag, chromatin accessibility (RNAPII-S5P CUTAC), and transcriptional activity (START-seq) at the Nanog promoter and enhancer cluster and flanking genes. Previously annotated enhancer regions^85^ are shown on top. **b, c,** Heatmaps (bottom) and average plots (top) comparing RNAPII, BRG1, and histone PTM occupancy by CUT&Tag, relative to the primary peaks (summits) of RNAPII-S5P CUTAC (b) and RefSeq annotated gene TSSs (c), sorted by decreasing RNAPII-S5P occupancy. **d,** Violin plots of CUT&Tag signal distribution comparing RNAPII-S5P and BRG1 occupancy at gene promoters (TSS) versus promoter-distal S5P CUTAC peaks (Distal). Median value (solid line), upper and lower quartiles (broken lines) and outliers were calculated using the Tukey method. **e,** Scatterplots comparing BRG1 and RNAPII S5P, S2P, and RPB3 CUT&Tag reads in 1000 bp genome-wide consecutive non-overlapping bins. **f,** Heatmaps (bottom) and average plots (top) comparing chromatin accessibility (RNAPII-S5P CUTAC and ATAC-seq), nucleosome positions (MNase seq), and transcriptional activity (START-seq), relative to the primary peaks (summits) of RNAPII-S5P CUTAC; and RNAPII-S5P CUTAC signal relative to RefSeq annotated gene TSSs (extreme right). Datasets are representative of at least two biological replicates.

**Extended Data Fig. 2.**
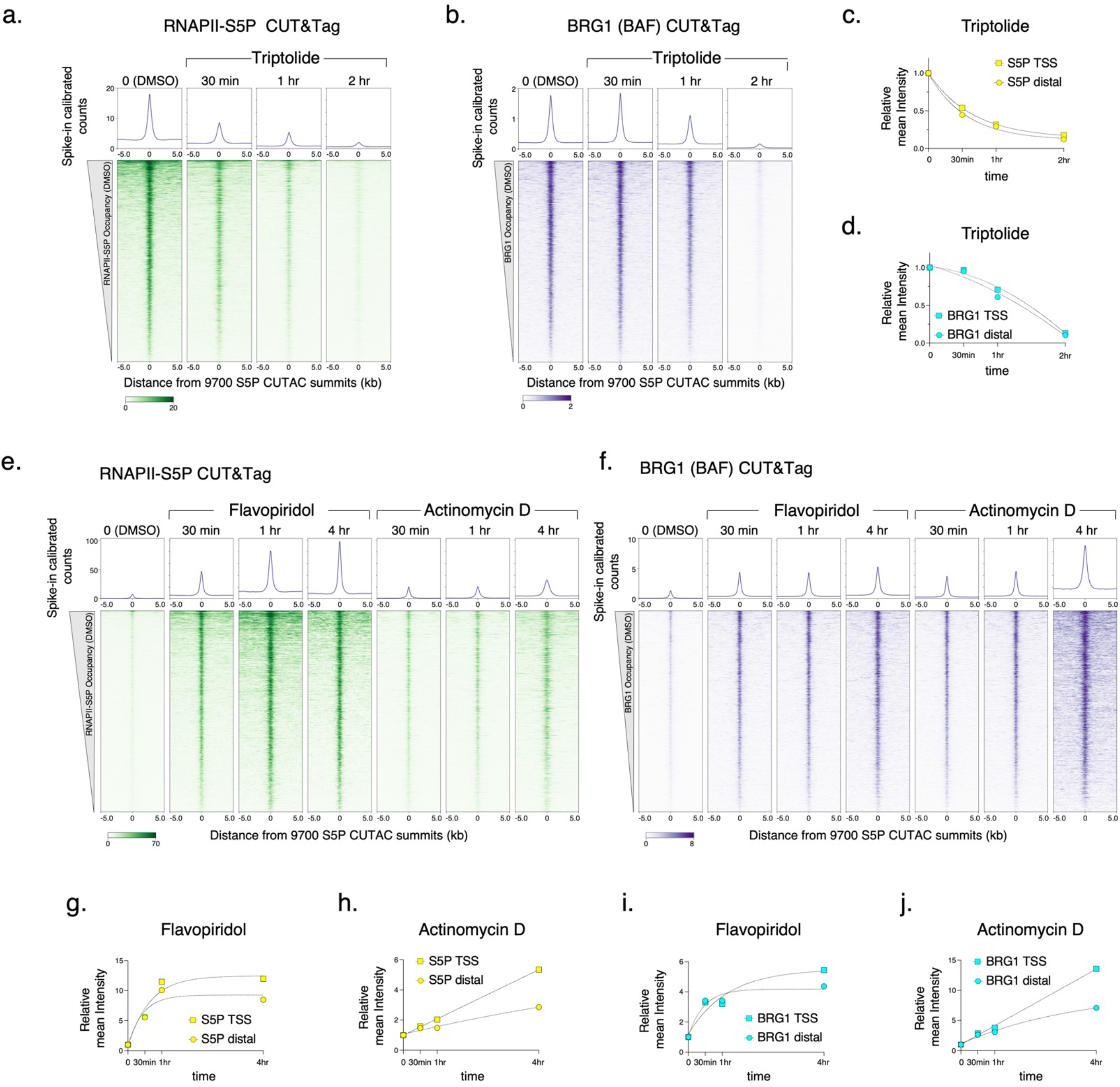
CUT&Tag of RNAPII-S5P and BRG1 after inhibitor treatment. **a, b, e, f,** Heatmaps (bottom) and average plots (top) comparing RNAPII-S5P and BRG1 occupancy by spike-in calibrated CUT&Tag relative to the primary peaks (summits) of RNAPII-S5P CUTAC in untreated cells (DMSO) versus cells treated with Triptolide (a, b), Flavopiridol, and Actinomycin D (e, f) at indicated time points post drug treatment. **c, d, g-j,** Comparison of fold changes in mean RNAPII-S5P and BRG1 occupancy (spike-in calibrated CUT&Tag) at gene promoters (TSS, squares) and promoter-distal CUTAC peaks (Distal, circles) at time points after drug treatments. Datasets are representative of at least two biological replicates.

**Extended Data Fig. 3.**
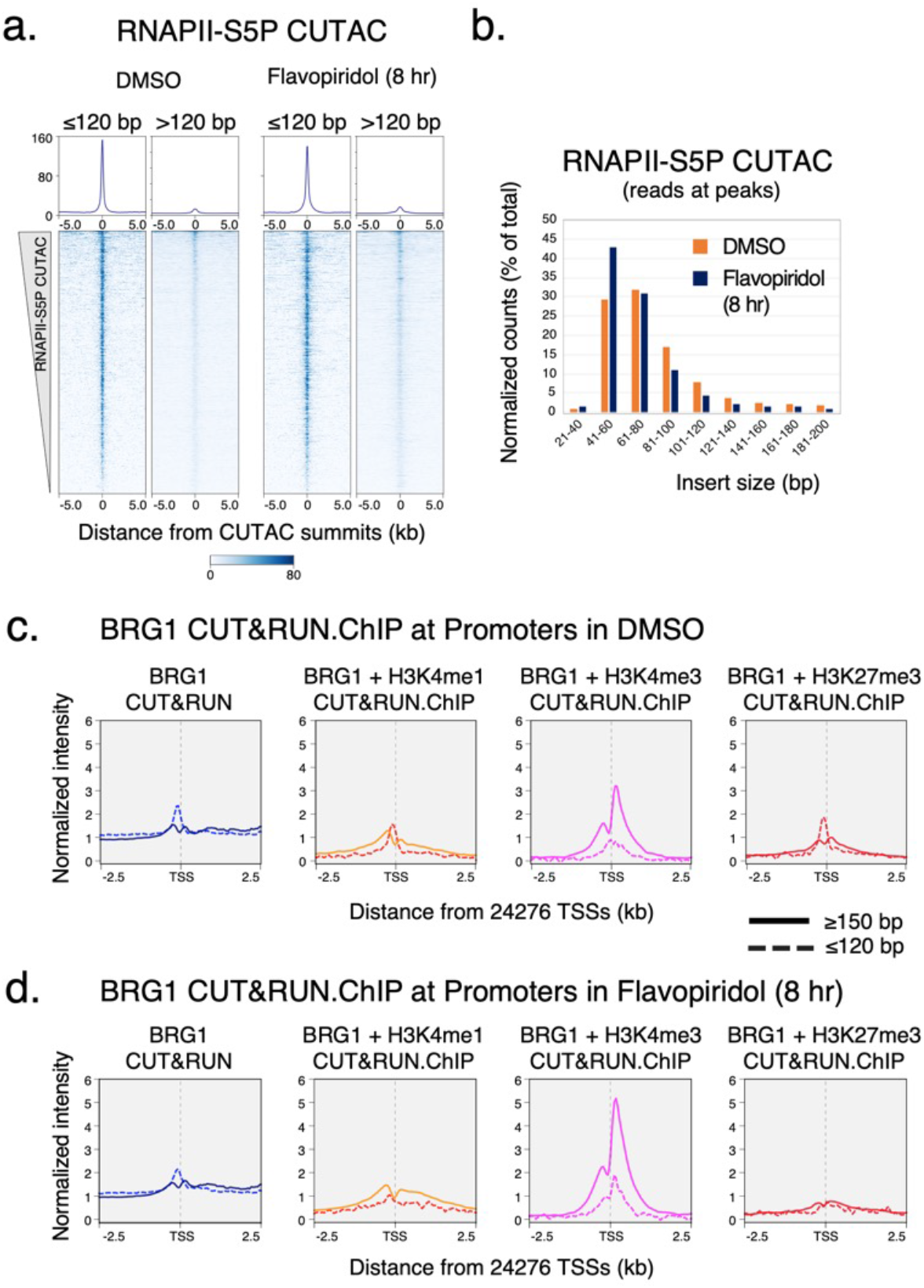
CUT&RUN.ChIP of BRG1 at promoters. **a,** Heatmaps (bottom) and average plots (top) of RNAPII-S5P CUTAC separated by fragment size, relative to primary peaks (summits) of RNAPII-S5P CUTAC. **b,** Comparison of RNAPII-S5P CUTAC fragment size distribution over peaks (promoter and enhancer NDR spaces) in cells treated with DMSO (control) and Flavopiridol; same data as used for Fig. 2a. **c, d,** Enrichment of nucleosomal (≥150 bp, solid lines) and subnucleosomal (≤ 120 bp, broken lines) reads from BRG1 CUT&RUN and CUT&RUN.ChIP experiments, relative to gene promoter TSSs, in DMSO and Flavopiridol treated cells. CUT&RUN.ChIP data were plotted as enrichment in histone ChIP over IgG isotype control. Datasets are representative of at least two biological replicates.

**Extended Data Fig. 4.**
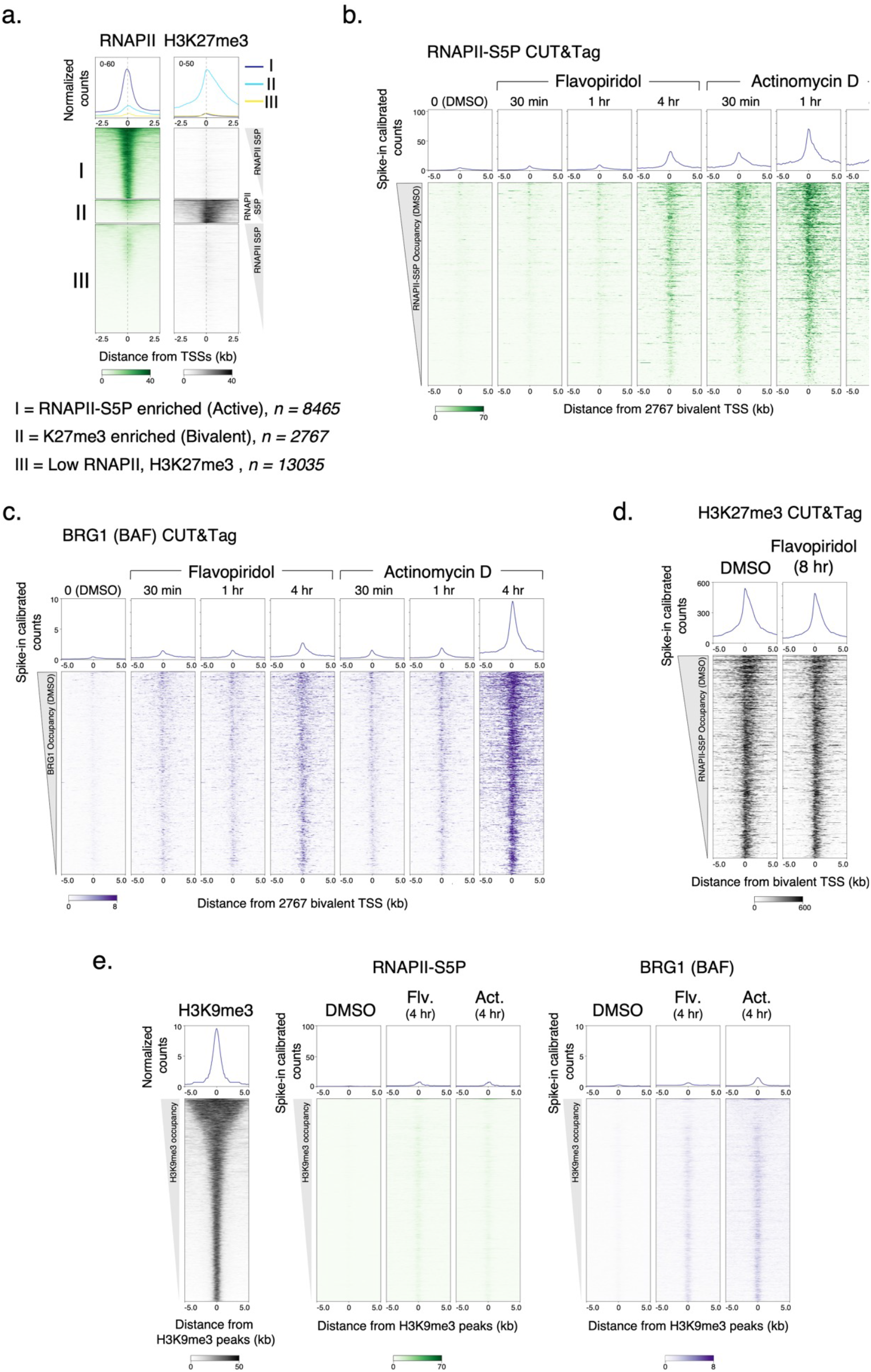
CUT&Tag of RNAPIIS5P and BRG1 at bivalent promoters. **a,** K-means clustering of RNAPII-S5P and H3K27me3 CUT&Tag reads relative to RefSeq annotated gene promoter TSSs to group promoters as active (I, RNAPII-S5P enriched) and bivalent (II, H3K27me3 enriched), and not enriched for either (III). **b, c,** Heatmaps (bottom) and average plots (top) comparing RNAPII-S5P and BRG1 occupancy by spike-in calibrated CUT&Tag relative to bivalent promoter TSSs in untreated cells (DMSO) versus cells treated with Flavopiridol or Actinomycin D at indicated time points post drug treatment. **d,** Heatmaps (bottom) and average plots (top) comparing H3K27me3 occupancy by spike-in calibrated CUT&Tag relative to bivalent promoter TSSs in untreated cells (DMSO) and cells treated with Flavopiridol. **e,** Heatmaps (bottom) and average plots (top) comparing H3K9me3 occupancy (CUT&Tag) with RNAPII-S5P and BRG1 relative to H3K9me3 peaks in untreated cells (DMSO) versus cells treated with Flavopiridol or Actinomycin D. RNAPII-S5P and BRG1 CUT&Tag reads were spike-in calibrated and plotted to the same scales as in panels b and c, respectively. Datasets are representative of at least two biological replicates.

**Extended Data Fig. 5.**
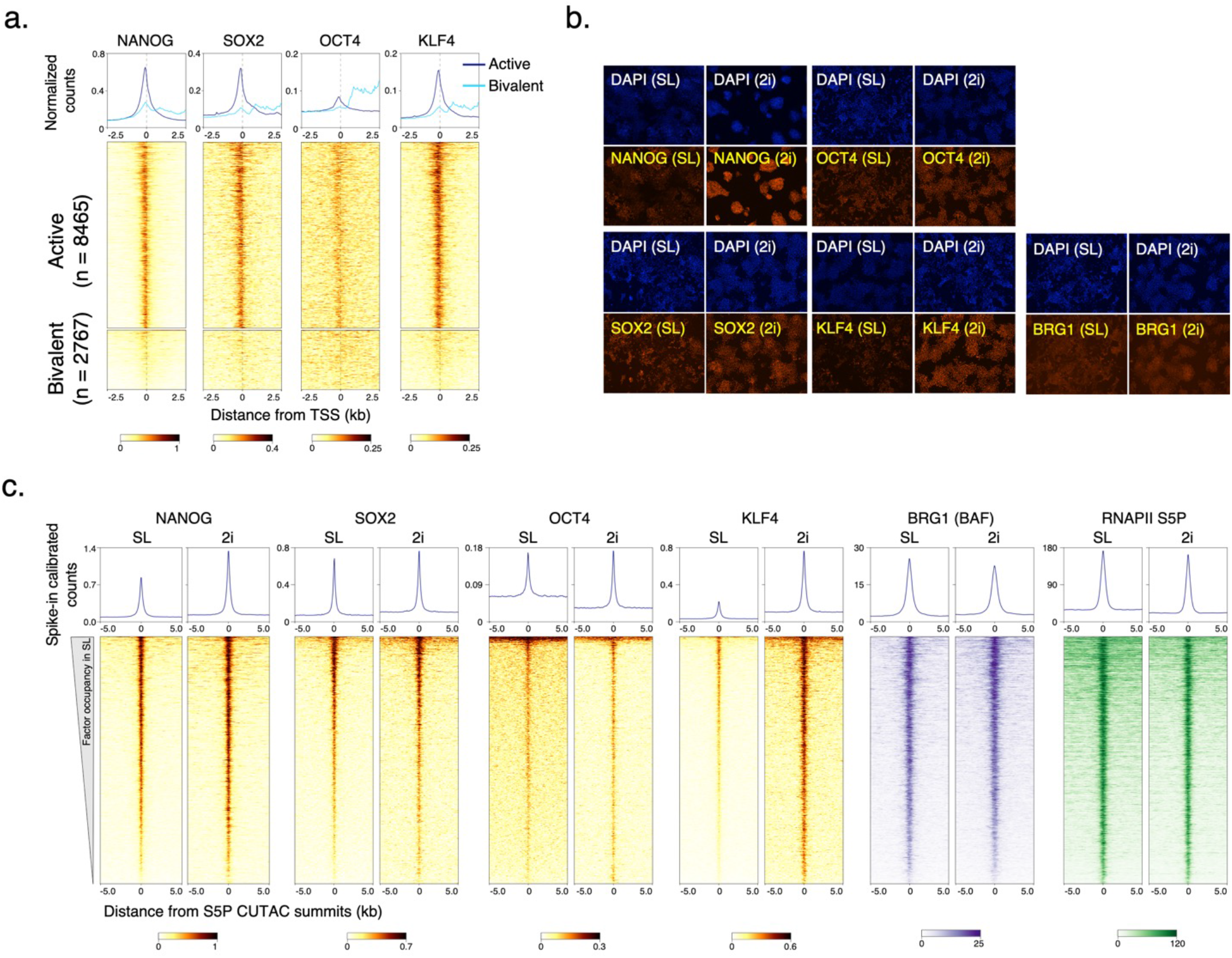
CUT&RUN of pluripotency TFs in SL versus 2i culture conditions. **a,** Heatmaps (bottom) and average plots (top) comparing pluripotency TF occupancy by CUT&RUN at RNAPII-enriched (active) and H3K27me3-enriched (bivalent) promoters. Promoters were grouped based on K-means clustering of RNAPII-S5P and H3K27me3 CUT&Tag reads mapping to a 5 kb window around the TSSs of RefSeq-annotated mESC genes, see Extended Data Fig. 4a. **b,** Immunofluorescent staining comparing pluripotency TF and BRG1 expression in SL versus 2i culture conditions. Cy5-conjugated secondary antibodies were used in all experiment except for KLF4, where Rhodamine red-conjugated antibody was used. DAPI (blue) was used to stain the nucleus in cells. **c,** Heatmaps (bottom) and average plots (top) comparing pluripotency TF occupancy by spike-in calibrated CUT&RUN in SL versus 2i culture conditions.

